# SUBTLE: An unsupervised platform with temporal link embedding that maps animal behavior

**DOI:** 10.1101/2023.04.12.536531

**Authors:** Jea Kwon, Sunpil Kim, Dong-Kyum Kim, Jinhyeong Joo, SoHyung Kim, Meeyoung Cha, C. Justin Lee

## Abstract

While huge strides have recently been made in language-based machine learning, the ability of artificial systems to comprehend the sequences that comprise animal behavior has been lagging behind. In contrast, humans instinctively recognize behaviors by finding similarities in behavioral sequences. Here, we develop an unsupervised behavior-mapping framework, SUBTLE (spectrogram-UMAP-based temporal-link embedding), to capture comparable behavioral repertoires from 3D action skeletons. To find the best embedding method, we devise a temporal proximity index as a metric to gauge temporal representation in the behavioral embedding space. The method achieves the best performance compared to current embedding strategies. Its spectrogram-based UMAP clustering not only identifies subtle inter-group differences but also matches human-annotated labels. SUBTLE framework automates the tasks of both identifying behavioral repertoires like walking, grooming, standing, and rearing, and profiling individual behavior signatures like subtle inter-group differences by age. SUBTLE highlights the importance of temporal representation in the behavioral embedding space for human-like behavioral categorization.

**One Sentence Summary:** Unsupervised behavior-mapping from 3D action skeletons achieves superior performance, captures behavioral repertoires, and identifies inter-group differences, emphasizing how temporal representation is critical in the behavioral embedding space.

## Main Text

Behavior categorization is the process of grouping different behavioral patterns into meaningful categories based on their characteristics (*1, 2*). Such categorization is a key learning attribute of the human brain and involves identifying common features that are shared across different behaviors while also creating meaningful distinctions between behavioral categories. This process is essential to how we understand and interact with the environment around us. Humans can categorize complex behaviors intuitively by observing only a few coherently moving key points (*3*). In contrast, it is far more difficult for machines to categorize the complex behavior of moving humans and animals, despite recent advances that enable machines to recognize 3D points accurately (*4-8*). Since we have yet to understand how humans intuitively recognize behavioral similarities, enabling machines to understand natural behavior by modeling human learning remains as a challenge.

Recent studies have tried to establish a machine-learning framework for categorizing behavior (*9-12*), particularly with supervised behavior mapping that uses human-annotated labels (*10, 11*). However, this approach is labor-intensive and prone to observer bias. Indeed, the process of manually annotating behavioral patterns can be time-consuming (minimum 3-4x the video’s duration varying performance across days), especially for large datasets, and the results may be influenced by the subjective judgments of the annotators(*10, 13*). As a result, researchers have sought unbiased approaches including unsupervised learning methods that do not require labor-intensive annotation by humans (*9, 10, 14-16*).

Unsupervised methods are, however, currently an ill-posed problem, as they cannot directly access labeled target outputs. Instead, incorporation of domain knowledge or additional constraints is required to guide unsupervised algorithms toward a more meaningful solution. In most cases, such methods aim to understand the underlying structure of the data to extract meaningful patterns and relationships. This is achieved through feature extraction, dimensionality reduction, and clustering.

Such data representation requires a robust feature embedding space to ensure the reliability and validity of the results. An embedding space allows relevant features and relationships between entities to be represented so that similar entities are close together and dissimilar entities are far apart (*17*). As behavior is a complex and dynamic process that changes over time, it is challenging to quantify its temporal structure without a clear ground truth (*2*). Therefore, unsupervised behavior mapping may require effective temporal representation.

While numerous approaches have aimed to construct an embedding space for behaviors, Berman *et al*. made a significant contribution by generating maps of fly behavior based on unsupervised approach (*14, 18*). They successfully assigned behavioral features to a low-dimensional embedding space using wavelet spectrograms for feature engineering and t-stochastic neighbor embedding (t-SNE) for nonlinear dimension reduction (*14*). From this embedding space, they employed predictive information bottleneck strategies to identify biologically meaningful repertoires (*18*).

While Berman *et al*. significantly influenced later studies and led to the emergence of other variants (*5, 7, 9, 12, 18-24*), the field has not defined criteria or reached a consensus on what constitutes a good behavioral embedding space (table. S1). Importantly, how the embedding space is designed is critical for predicting meaningful and reliable categories. In the absence of a standard evaluation method, however, previous research has yet to evaluate the quality of embedding spaces and has instead focused on optimizing the downstream task.

Humans can learn new concepts from just a few samples by leveraging prior knowledge and using probabilistic reasoning to build new concepts from the observed examples. This process is thought to involve the utilization of temporal information and transition probabilities for sequence coding, which allows us to represent and utilize the probabilities of future events and the time intervals between those events to guide our choices and behavior. The human brain constructs categorical representations, potentially serving as higher-level feature embeddings (*1, 25, 26*), and tracks the transition probability between events to anticipate future events, exhibiting a stronger response to unexpected events compared to expected events.

Introducing human-based constraints into machine learning models may improve performance, as research has shown that machines can achieve human-level concept learning through probabilistic program induction (*27*) and natural language description (*28, 29*). The process by which humans categorize behaviors is not fully understood, but it is known that the brain uses temporal dynamic information encoded in embedding space for categorization. For example, a temporal choice task has been utilized to investigate temporal and probabilistic calculations in both mice and humans (28, 29). As we further investigate the human brain’s ability to process and utilize temporal information and transition probabilities (*30*), we can potentially enhance machine learning models to mimic human learning capabilities, bridging the gap between human and machine cognition.

To enable machines to categorize behaviors in a manner similar to that of humans, we have developed a simple metric to measure temporal representation in behavioral embedding space. To evaluate the quality of the embedding space, we assume that a good embedding space should preserve the temporal representation of behaviors, which is reflected in the distances between embeddings (Fig. 1A and B). Based on this premise, we have developed an evaluation method to quantify the temporal connectivity (Fig. 1C and D) by calculating the transition probability between two clusters in the space. We consider two cluster centers to be temporally linked in the embedding space if they are in close proximity and have a high transition probability. Given the number of clusters (*n*) and their centroids {*c*_1_,…,*c*_*n*_} we define the temporal proximity index (TPI) for quantifying temporal representation as the following equation:

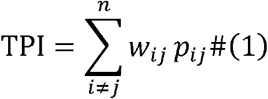

*p*_*ij*_ is a transition probability from source cluster *c*_*i*_ to target cluster *c*_*j*_. *w*_*ij*_ is a proximity measure, which is defined as

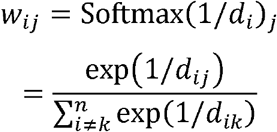

*w*_*ij*_ is calculated from source to target cluster distances *d*_*i*_ = {*d*_*ij,j* ≠*i*_ | *j* = 1, …, *N* }where *d*_*ij*_ = |*c*_*i*_ − *c*_*j*_|_2_, an *L*_2_-norm or Euclidean distance between two clusters. Thus, *w*_*ij*_ represents the relative proximity between the source cluster *i* and the target cluster *j*, measured by Softmax of the inverse distance 1/*d*_*ij*_. Overall, TPI can be interpreted as the sum of proximity-weighted transition probabilities. To implement the concept of TPI, we designed a workflow similar to the previously described unsupervised behavior mapping framework by creating feature embeddings, subclusters, and superclusters (Fig. 1D) (*14, 18*).

**Fig. 1.**
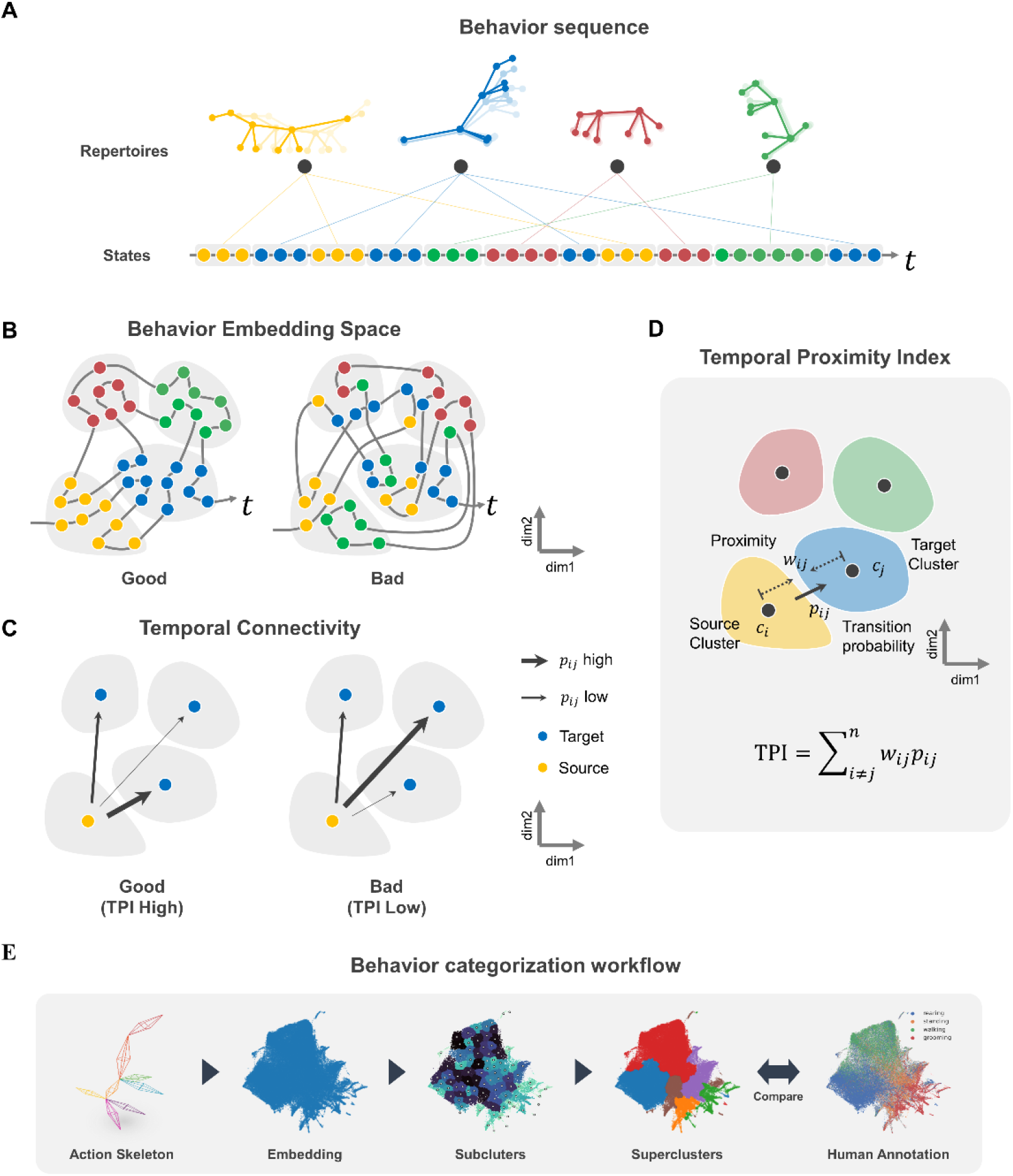
Evaluation strategy for temporal representation in the behavioral embedding space. (**A**) Conceptual illustration of an example behavior sequence for the 3D action skeleton trajectory of a mouse, where each color represents stereotyped behavioral repertoires. (**B**) Illustration of example pattern trajectories in the behavior embedding space, indicating embedding quality can determine more efficient trajectories; left, good embedding space with each cluster containing the same behavioral states with efficient temporal trajectories; right, bad embedding space with each cluster contains the different behavioral states with inefficient temporal trajectories (**C**) Illustration of example temporal connectivity in the behavior embedding space represented with transition probabilities from source to target clusters; left, good temporal connectivity with closer distances between clusters as the transition probability increases; right, bad temporal connectivity with greater distances between clusters as the transition probability increases. (**D**) Temporal Proximity Index (TPI), for assessing the temporal connectivity of the behavior embedding space. (**E**) Schematic workflow for unsupervised behavior categorization and evaluation in this study.

To create feature embedding from action skeletons, we obtained 3D action skeleton trajectories from freely moving mice (N = 10, ≈120,000 video frames) using the AVATAR system (Fig. 2A, left) (*8*). The system helps identify target behavioral categories from logs such as walking, grooming, rearing, and standing. For this, the system (Fig. 2A, left) utilizes five cameras to record multi-view images, while AVATARnet captures key points via object detection and reconstructs their trajectories in 3D Euclidean space (Fig. 1). As part of the input features (Fig. 2A, right top), we extracted additional information from the Morlet wavelet spectrograms and kinematic features from the raw coordinates of 3D action skeleton trajectories. For the nonlinear mapping (Fig. 2A, right bottom), we applied t-stochastic neighbor embedding (t-SNE) (*31*) and uniform manifold approximation and projection (UMAP) (*32*) on the three input features to generate six combinations of feature embeddings. Two examples of feature embeddings are shown in Fig. 2B.

**Fig. 2.**
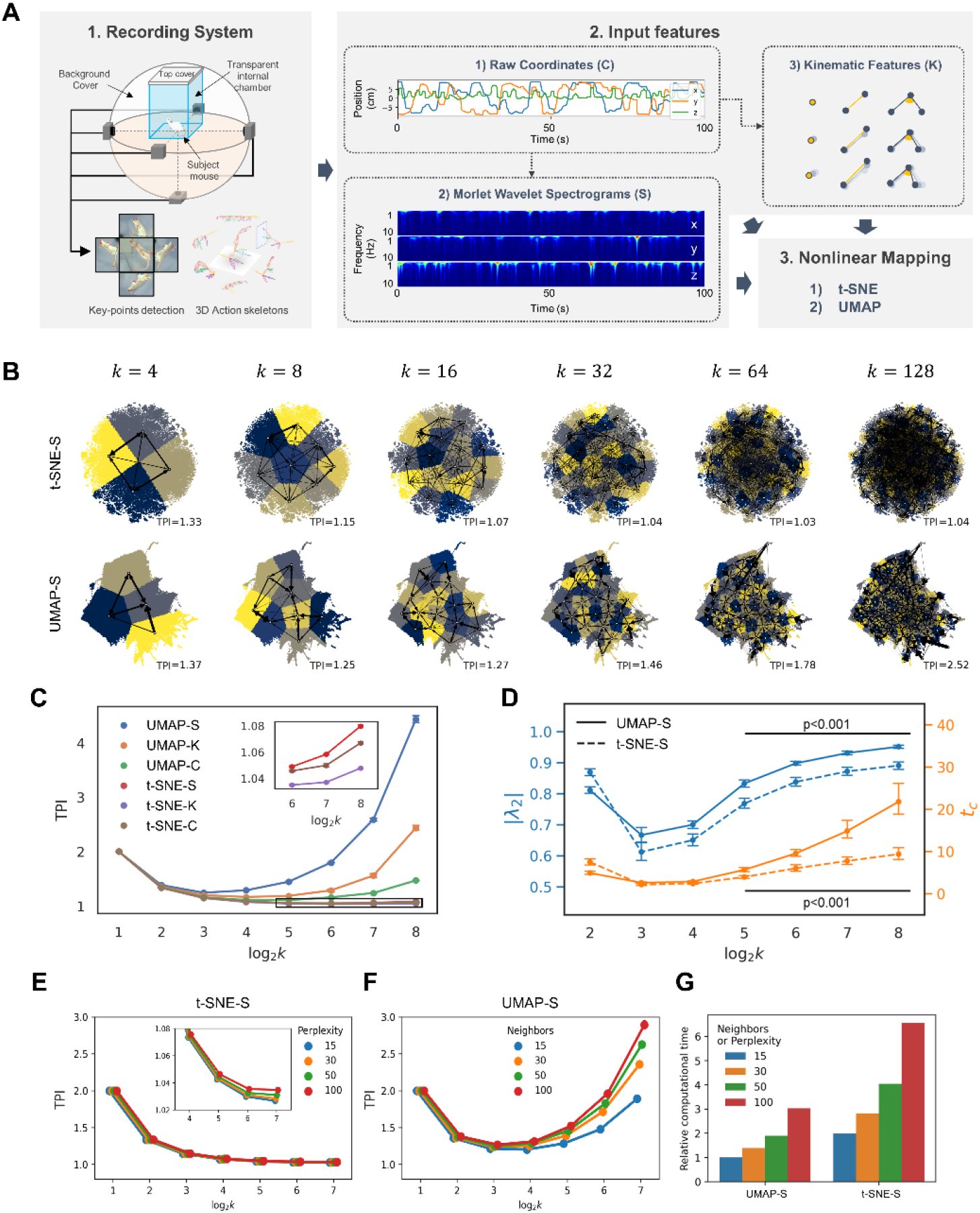
SUBTLE method gives the highest temporal connectivity score in embedding space. (**A**) Schematic flowchart for preparing various feature embedding methods. 1) 3D action skeletons were extracted with the AVATAR recording system. 2) Kinematic features and wavelet spectrograms were extracted from raw coordinates. 3) Nonlinear mapping with t-SNE and UMAP algorithms. (**B**) Inter-cluster transition probabilities are visualized on embedding spaces as the number of clusters of *k*-Means clustering increases. (**C**) TPI scores were calculated with three input features (coordinates, kinematics, spectrogram) and two nonlinear embedding methods (t-SNE, UMAP). Inset shows the magnified plot of the black box containing t-SNE results. (**D**) Comparison of the second largest eigenvalue λ_2_ and its characteristic time t_2_ of the transition matrix obtained from spectrogram-UMAP and spectrogram-t-SNE with respect to increasing cluster numbers in *k*-Means clustering. (**E**) Effect of increasing perplexity in t-SNE algorithm. Inset, the magnified plot from log_2_ k = 4, 5, 6, 7. (**F**) Effect of increasing neighbor in UMAP algorithm. (**G**) Comparison of computational efficiency between spectrogram-UMAP and spectrogram-t-SNE with respect to Neighbors or Perplexity hyperparameters (15, 30, 50, 100).

Next, we evaluated the temporal representation of each embedding space using TPI (Eqn. 1). By conducting *k*-Means clustering with increasing cluster numbers, *k*, we could enhance the resolution of temporal connectivity (Fig. 2B). While the spectrogram-UMAP feature embedding showed a significant improvement in TPI scores at high *k*, spectrogram-t-SNE did not (Fig. 2B). We found that using the wavelet spectrogram in combination with UMAP resulted in the highest TPI value or the best temporal representation, outperforming the other five combinations (Fig. 2C). These results indicate that the wavelet spectrogram reflects the temporal representation of action skeleton trajectories more accurately than raw coordinates or kinematic features (Fig. 2C).

When it comes to the nonlinear mapping, we found that UMAP has higher TPI values than t-SNE (Fig. 2B and C), possibly due to its ability to preserve the global structure (*33*). To test this possibility, we systematically altered the global structure of embedding space by changing the main hyperparameters for t-SNE (with perplexity) and UMAP (with neighbors). Regrettably, there is no universally optimal number of hyperparameters for neighbors in UMAP and perplexity in t-SNE, general recommendation ranges from 15 to 100 (common default values for neighbors=15 and perplexity=30). Generally, increasing the values of these hyperparameters will prioritize the preservation of global structure in the embedding space at the expense of local structure. While we observed a general increase in TPI scores for both t-SNE and UMAP by increasing these hyperparameters, the increase was more pronounced for UMAP (Fig. 2E and F), implying that the ability of UMAP to better preserve global structure compared to t-SNE contributes to enhancing temporal connectivity in the embedding space. In addition, UMAP was computationally efficient compared to t-SNE, highlighting its practical advantage (Fig. 2G). Balancing computational cost and TPI values, we set the UMAP neighbors parameter to 50, optimizing the preservation of global structure and efficiency in analyzing temporal connectivity.

Next, to investigate the time scale of temporal dynamics in this embedding space, we measured the characteristic time (i.e., information decay of the behavioral state) of state transitions following a previous approach (*18*). We consider the number of clusters, *k*, to be behavioral states, and if the dynamics of the behavioral states follow Markovian dynamics, the transition from one state to another is independent of history. However, the Markov model has almost no information beyond 40 steps (2 seconds), while the transition matrices from actual data for τ=40 (2 seconds) and τ=400 (20 seconds) retain information (see fig. S5A). These results support the notion that the dynamics of freely moving mice are not Markovian, and they exhibit a much longer time (at least more than 2 seconds) structure than the Markovian model. In the Markovian system, the transition matrix **T** provides a complete description of the dynamics.The slowest time scale in the Markovian system is determined by the second largest eigenvalue, | λ_2_|, which results in a characteristic decay time (the average time point at which 1/e of information is lost), *t*_*c*_ = −1 / log|λ_2_| (*18*).We obtained the |λ_2_|, and *t*_*c*_ from **T** constructed from *k* number of clusters for spectrogram-UMAP and spectrogram-t-SNE. We found that spectrogram-UMAP gives a higher characteristic time than the spectrogram-t-SNE at log_2_ *k* > 4 (Fig. 2D). This result implies that information on behavioral states decays significantly more slowly in spectrogram-UMAP than in spectrogram-t-SNE. Based on these results, we conclude that the spectrogram-UMAP-based temporal-link embedding (SUBTLE) offers the better preservation of temporal representation in the behavioral embedding space.

As kinematic features provide a quantitative description of how an organism moves, we hypothesized that a good behavioral embedding space should manifest kinematic characteristics. We assessed the kinematic similarity in the embedding space by calculating the inter-embedding cosine similarity and relative inter-embedding distance of kinematic features (Fig. 3A). The results indicate that the SUBTLE effectively reflects kinematic similarity as a distance, in contrast to the t-SNE-based approach (Fig. 3B and C). Additionally, the visualization of the first principal component (PC1) of kinematic features in the embedding space highlights the effectiveness of the spectrogram-UMAP method (Fig. 3D). In summary, our proposed behavior embedding method, SUBTLE, preserves both temporal representation and kinematic structures and excels at organizing kinematic features.

**Fig. 3.**
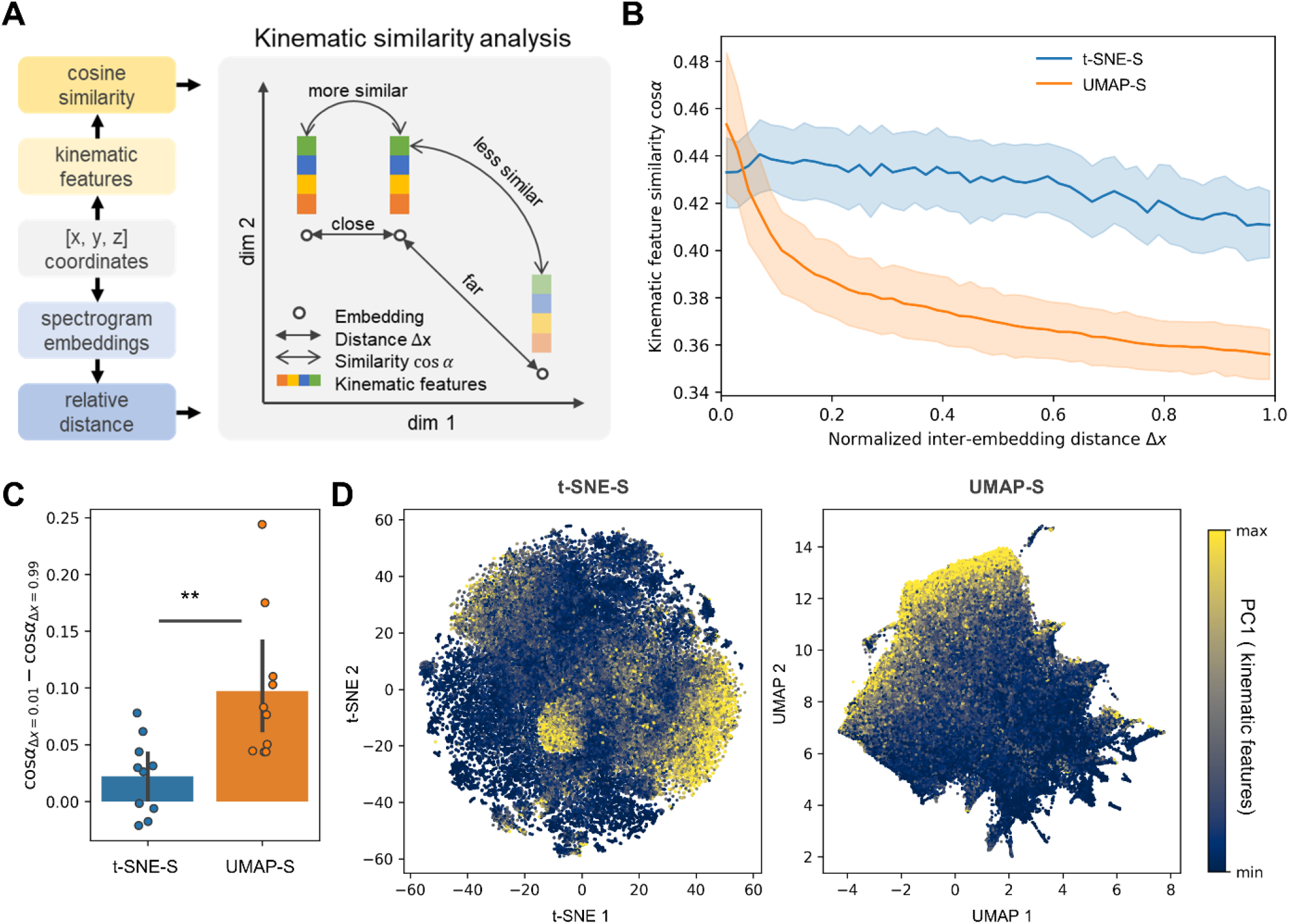
SUBTLE effectively reflects kinematic feature representation in embedding space. (**A**) Schematic illustration of kinematic similarity analysis in the behavioral state embedding space. Note that kinematic features were not directly included while constructing the embedding space. (**B**) The relationship between inter-embedding kinematic feature similarity cos_α_ and normalized embedding distance Δx. (**C**) Perspective contrast of inter-embedding kinematic similarity in mice (n=10). (**D**) The first principal component of kinematic features was plotted on the embedding spaces constructed from spectrogram-t-SNE and spectrogram-UMAP.

As a next step, we identified unique stereotyped actions (e.g., lifting, stretching, and balancing) as behavioral states (subclusters) (*14*) which can be used as building blocks for behavioral repertoires (superclusters) that reflect sequential transitions of behavioral states (e.g., rearing) (*18*). Several options are available to support this stage, including the Watershed algorithm (*14*), *k*-Means clustering (*34*), Gaussian Mixture (*35*), and HDBSCAN (*9*). However, these algorithms require prior knowledge or assumptions about the embedding space, such as a probability density function, a predetermined number of clusters, or a threshold density. To overcome this, we used Phenograph clustering (*36*), a graph clustering algorithm that can cluster data points directly using Jaccard similarity and automatically determine the optimal number of clusters by maximizing the modularity of the clusters. Phenograph clustering produced better inner cluster inter-embedding kinematic similarities than *k*-Means clustering (fig. S2), probably due to its ability to distinguish isolated groups in the embedding space (fig. S2A, yellow arrow). This demonstrates using the Phenograph clustering algorithm in SUBTLE lead to finding higher-quality clusters.

We then utilized these subclusters to build a transition matrix between behavioral states. The state transition probabilities (shown by arrows in Fig. 4A) and the state retention rate (depicted by a hue in Fig. 4B) were used to determine the biological significance of these values. When comparing the retention rate with human-annotated labels (Fig. 4C), grooming behavioral states were commonly found in clusters with high retention rates, whereas walking or rearing was frequently found in clusters with relatively low retention rates (depicted by color in Fig. 4G and fig. S4). While spectrogram-t-SNE gives similar patterns, the effect is more pronounced in the spectrogram-UMAP approach used in SUBTLE.

**Fig. 4.**
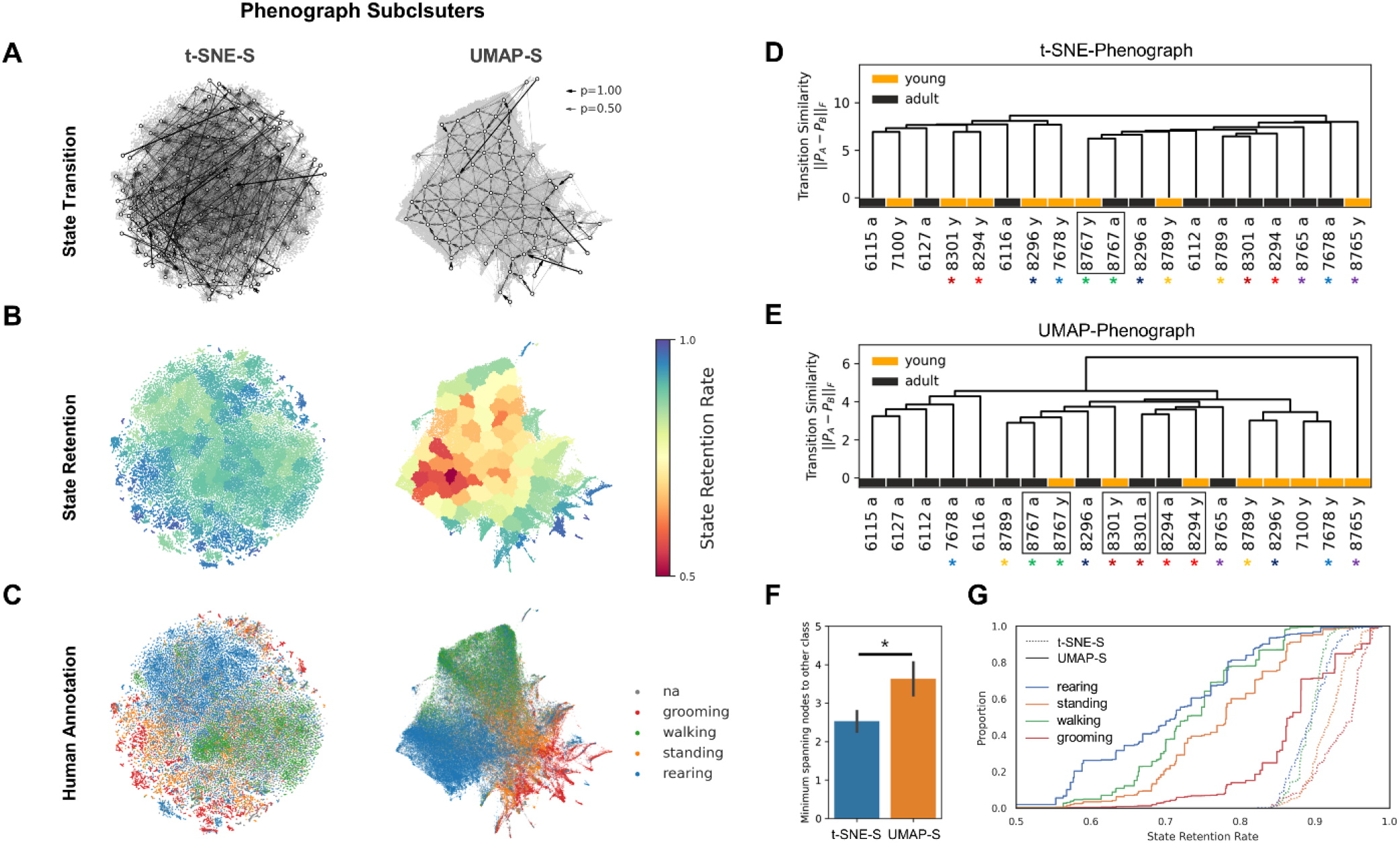
SUBTLE-based subclusters effectively capture biologically meaningful features. (**A**) State transition map of Phenograph subclusters in embeddings of spectrogram-t-SNE and spectrogram-UMAP (**B**) State retention map of Phenograph subclusters in embeddings of spectrogram-t-SNE and spectrogram-UMAP (**C**) Human-annotated behavior categories in spectrogram-t-SNE and spectrogram-UMAP (**D-E**) Dendrogram of inter-subject behavior transition similarity measured by Frobenius norm of two different transition probability matrices P_A_ and P_B_. Asterisks with the same colors denote the same subject with different recording periods, young and adult. (**F**) Comparison of minimum spanning nodes to join the other class in the dendrogram. (**G**) Empirical cumulative distribution function of human-annotated behavioral states against retention rate.

Next, we created a dendrogram using the similarity distance between animal subjects, calculated using the Frobenius norm of the behavior transition matrix (represented by color in Fig. 4D and E). The UMAP-based method distinguished young and adult mice better than the t-SNE method (Fig. 4F). Interestingly, data from the same subjects recorded in their young and old life stages (fig. S3) appear close to each other when age information was removed (Fig. 4E). This result suggests that the information about state transitions of the subclusters created from the spectrogram-UMAP-based embedding space can be used to extract biologically meaningful insights without the need for human-provided behavior labels. In conclusion, we have developed Phenograph clustering and spectrogram-UMAP-based embeddings that can identify and interpret behavioral patterns in embedding space, which help extract biologically meaningful insights without human annotation.

To obtain unsupervised behavior categories, we employed the predictive information-based deterministic information bottleneck (DIB) algorithm (*18, 37, 38*). The algorithm aims to identify common patterns of an animal’s behavioral repertoires (superclusters) for accurate predictions of future behavioral states (subclusters), while minimizing reliance on past behavioral states by finding the optimal trade-off between predictability of the future and reliance on past. We grouped the subclusters into *n* superclusters that contained information about the future state after τ steps while retaining less information about the past state (see Superclustering method section in Method for more details). As a result, we identified superclusters for n = 2,4,6 (Fig. 5A) and human-annotated labels have similar structures to the *n* = 6 case (see Fig. 5B). For *n* = 6, the transition matrix of subclusters with grouping by the obtained superclusters is shown in Fig. 5C. Clustering results and the grouped transition matrix for cases where *n* ranges from 1 to 7 can be found in fig. S5B and S6C. A human-interpretable hierarchical structures can be observed in the Sankey diagram flow (see Fig. 5D) as *n*, increases. An example of the 30-second animal behavior, an ethogram, is also presented for *n =* 2, *n =* 6, subcluster, human-annotated cases in Fig. 5E.

**Fig. 5.**
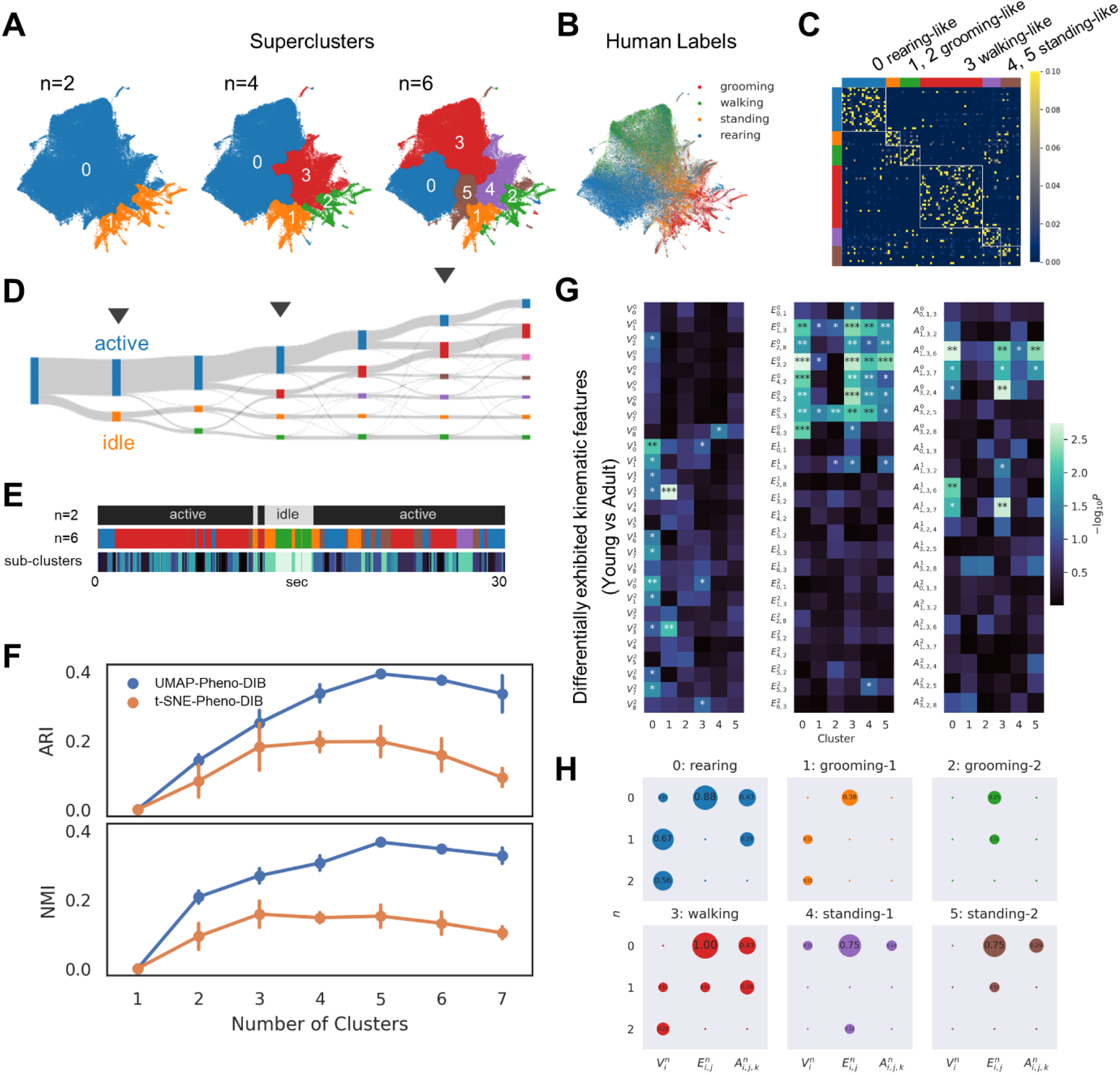
Comparison of SUBTLE-based superclusters and human-annotated behavior categories. (**A**) Hierarchical clustering with deterministic information bottleneck method, using lag τ = 2t_c_ ≈ 20. Identified representative superclusters (n = 2, 4 6) in UMAP (**B**) Human-annotated behavioral categories in UMAP (**C**) Example transition probability matrix of subclusters grouped by superclusters (n = 6). (**D**) Sankey diagram flowing horizontally from left to right with the increasing number of clusters (**E**) Example ethogram showing 30-seconds of mouse behavior (**F**) Comparison of superclusters with human-annotated behavioral categories using adjusted rand index (top) and normalized mutual information (bottom). (**G**) Differentially exhibited kinematic features of young and adult mice. (**H**) Proportion of significantly different features in each behavioral cluster.

We assessed the accuracy of the superclusters from the DIB algorithm compared to human-annotated animal behavior labels (Fig. 5F) using Normalized Mutual Information (NMI) and the Adjusted Rand Index (ARI), two widely used metrics for evaluating the performance of unsupervised clustering algorithms. We found that as the number of superclusters (*n*) increases, the clustering structure of the UMAP embedding space becomes more similar to the human-annotated behavior classification compared to t-SNE for all values of *n* that are available (see Fig. 5F and fig S6). We also confirmed the actual clustering result by viewing a recorded video, 3D action skeleton, human annotation, and SUBTLE supercluster simultaneously (fig. S7).

To assess age-dependent behavioral changes, we identified differential kinematic features found in superclusters of young vs. adult mice (Fig. 5E). We found most of the significant differences from zero derivatives of edges in all 6 superclusters, which may be due to the varying size between young and adult mice (Fig. 5E). Except for zero derivatives of edges, the majority of differences were found in rearing (17 features) and walking (9 features) clusters (Fig. 5E), indicating that young and adult mice have substantial differences in terms of dynamic movement. The smallest difference between young and adult mice was found in grooming-like behaviors (supercluster 1 and 2). The proportion of significantly different features is distinct for each superclusters (Fig. 5F). While our differential analysis of inter-group kinematic features provides interpretable enriched data, conventional approach that estimates locomotor activity by 2D-trajectory distances in open-field like environment revealed no significant inter-group difference (fig. S3C and D). In conclusion, by combining superclusters and kinematic feature analysis, our method successfully distinguishes subtle behavioral changes between young and adult mice.

We investigated the effect of random shuffles on the sequence of behavioral states to better understand the role of temporal connectivity in behavioral categorization. Specifically, we divided the sequence of behavioral states into equal-sized chunks and randomly swapped the order of the chunks to create a shuffled sequence (see Fig. 6A for an illustration with chuck size = 4). We then measured the effect of shuffling on the temporal connectivity of the data using the TPI. We found that as the chunk size decreased, the TPI metric was disrupted, indicating no temporal connectivity in the shuffled data (see Fig. 6B). Additionally, we evaluated the effect of shuffling on the performance of the superclustering algorithm using ARI and NMI scores. We found that as the chunk size decreased, ARI and NMI scores were also disrupted (see Fig. 6C). These results indicate that our TPI metric effectively captures temporal dynamics information and underscores the importance of temporal connectivity in behavioral categorization.

**Fig. 6.**
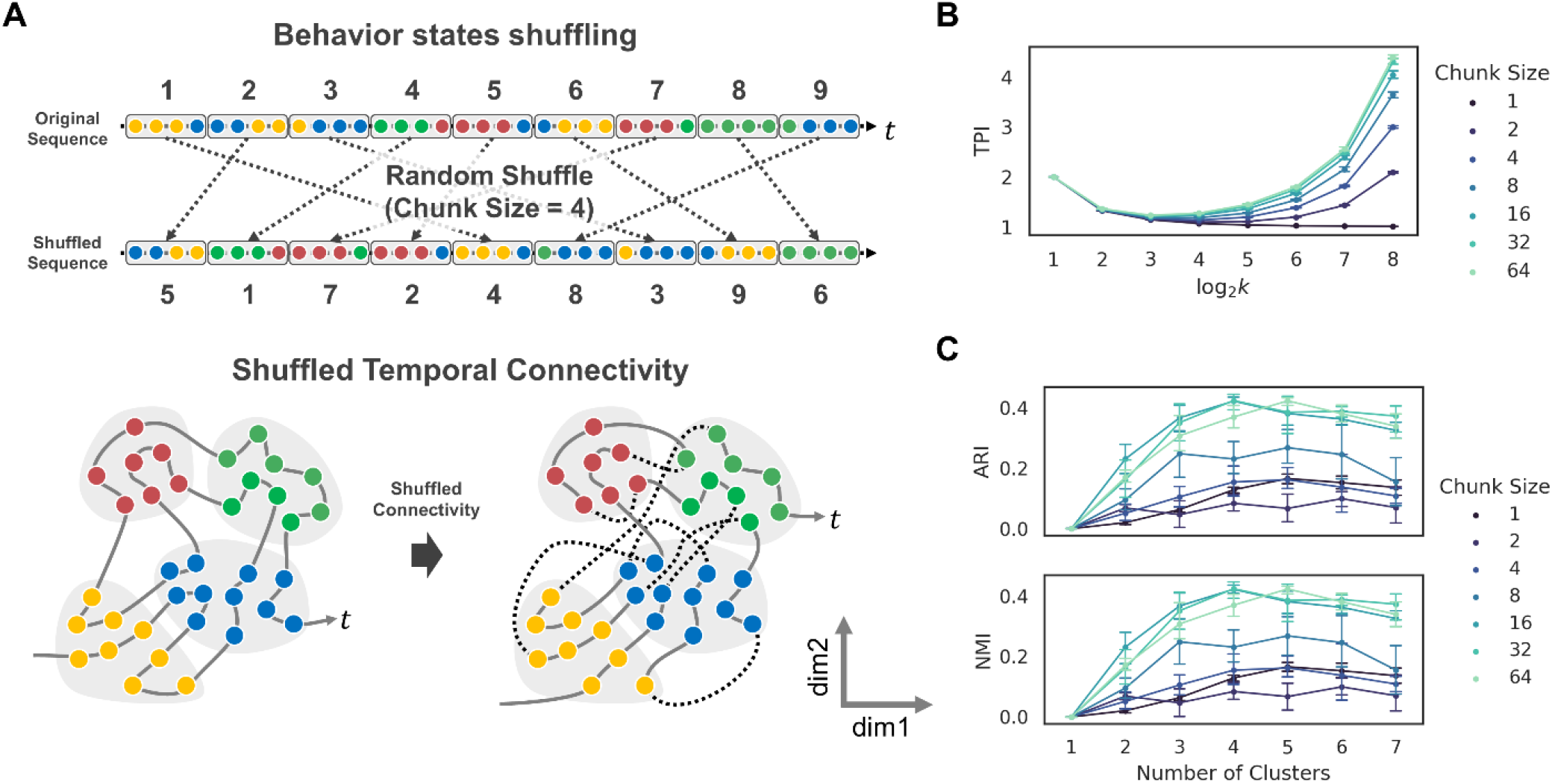
Importance of temporal sequence in human-like behavior categorization. (**A**) Conceptual illustration of shuffling the behavioral states with fixed chunk size (top) and shuffled temporal connectivity in behavioral embedding space (bottom). (**B**) Evaluation of temporal representation with TPI metric in shuffled embedding space with varying numbers of chunk sizes (**C**) Evaluation of clustering quality of superclusters using the shuffled embedding with varying numbers of chunk sizes.

## Discussion

Despite the success of deep learning in detecting 3D points in behaving animals, accurately identifying higher-order behavioral patterns remains a challenge for machine learning systems. While humans can contribute and classify animal behaviors using their intuition and labeling guidelines, incorporating human annotation into artificial behavioral analysis is labor-intensive and prone to observer bias. This limitation exists especially for large datasets, and the results may be influenced by the subjective judgments of the annotators. As a result, researchers have sought alternative approaches to overcome these roadblocks, including unsupervised methods that do not require human-provided labels. However, this approach raises a dilemma: the outcomes may differ from those of humans as the model does not have constraining or intuitive guidelines.

In this study, we developed a novel method called SUBTLE (spectrogram-UMAP-based temporal-link embedding) to map and analyze animal behavior. We evaluated the quality of behavioral embedding space by measuring the temporal representation using the proximal transition index (TPI). Our results show that using the wavelet spectrogram in combination with UMAP results gave the best temporal connectivity, outperforming other feature embedding methods. Additionally, SUBTLE effectively reflects kinematic similarity as a distance more effectively than other approaches. Our findings suggest that SUBTLE can help analyze animal behavior, as demonstrated in the analysis of ∼190 minutes (∼228,000 frames, 19 mice) long logs of mice movements and the corresponding ∼100 minutes (∼120,000 frames, 10 mice) human annotations. Furthermore, our study highlights the importance of temporal dynamics information in behavioral analyses and the value of using feature embeddings to extract meaningful information from raw data.

One of the key contributions of our study is the suggestion of an evaluation strategy, including the development of the TPI metric to assess temporal connectivity in the feature embedding space. This allows for a more rigorous evaluation of the method and provides insight into how to construct a behavioral embedding space that enables machines to categorize animal behavior similar to humans. Moreover, the proposed hypothesis on human brain using transition probabilities for sequence categorization well matches the concept of behavioral state transitions in the SUBTLE framework. Our proposed TPI metric and kinematic similarity measurement can be used for explicit constraints for setting learning objectives in AI-based modeling. The recent successes of large language models in natural language processing suggest the exciting possibility of creating large behavior models that could advance our understanding of animal behavior. These possibilities await future investigation by researchers, and we look forward to the potential breakthroughs that may arise from these developments.

Our SUBTLE framework offers several advantages for researchers. Firstly, it saves a significant amount of time by automating the identification of behavior repertoires, eliminating the need for labor-intensive manual annotation. Secondly, it is free from human subjective bias, resulting in a more reliable and unbiased identification of behavior repertoires. Thirdly, the framework allows for the identification of individual behavior signatures, which can be used for comparison with other individuals or groups (Fig. 4D-F). Finally, it can profile inter-group subtle kinematic feature differences, providing valuable insights into the nuances of animal behavior (Fig. 5G and H). These advantages make SUBTLE a valuable tool for advancing our understanding of animal behavior and potentially saving time and resources for researchers. Our spectrogram-based unsupervised framework and mouse action skeleton data will be released to the public (see Data and materials availability in Acknowledgement).

## Materials and Methods

### Animals

Young (3 - 4 weeks) and adult (8 - 12 weeks) Balb/c male mice (IBS Research Solution Center) were used for the acquisition of action skeletons. The animal experimental procedures were approved by the Institute for Basic Science (IBS; Daejeon, Korea). The animals were kept on a 12 hr light-dark cycle with controlled temperature and humidity and had ad libitum access to food and water.

### Behavior recording procedure

Recordings of behavior were performed after 3 days of handling and habituation. In order to reduce background stress and anxiety in the laboratory mice, cotton gloves were used during the entire handling process. At least 2 hours before the experiment, animals were moved to the experimental room for habituation. Spontaneous animal movements were recorded using the AVATAR system (*8*) for 10 minutes as a single session on 5 consecutive days. Between each session, the inner transparent recording chamber was cleaned with 70 % ethanol and dried thoroughly. To prevent the effect of noise, the recording system was set up in a sound-attenuated room.

### Data acquisition

We collected action skeletons with AVATAR, a YOLO-based 3D pose estimation system with multi-view images that extracts continuous joint movement from freely moving mice (*8*). This system is commercially available at https://www.avataranalysis.com/. The recording hardware consists of an inner transparent chamber and an outer recording box. The outer recording box consists of 5 Full-HD CMOS cameras (FLIR, U3-23S3M/C-C, 1200 × 12000 pixels, 20 frames per second): four at the side and one at the bottom, with a distance of 20 cm (side), and 16 cm (bottom) from the inner transparent chamber. These cameras were directly connected to the recording computer (CPU, Intel, i9-10900K; GPU, Colorful, RTX3080; RAM, 16GB). As high-resolution image acquisition is required to detect an animal’s joint coordinates accurately, the recording was conducted under bright lighting conditions. The brightness of the lighting inside the recording box was 2950 lux. The inner transparent chamber was positioned at the center of the outer recording box and measured 30 cm × 20 cm × 20 cm (height × width × length). Numerous breathing holes with a diameter of 5 mm were made on the upper side of the chamber. The top of the transparent inner chamber is covered with a white background lid after the animal enters. Acquired video files were processed with a pre-trained YOLO-based DARKNET model to extract the coordinates of mouse joints. To reflect real-world scales, the grid-like black and white panels were used for calibration (see fig. S1 for detail).

### Kinematic features

Here we provide a general definition of kinematic features. A mouse action skeleton dataset is given as continuous frames of multiple joint *v*_*t*_ of 2D (x_t_, y_t_) or 3D (x_t,_ y_t,_ z_t_) coordinates, where t is the specific frame. Given *T* number of time sequences and joint coordinates *v* = {*v*_t_ | *t* = 1, …, *T*}, we define nodes, edges, and angles as following:

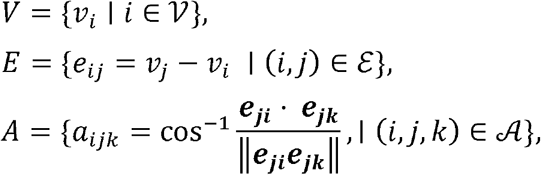

where 𝒱 is a set of joints, ℰ and 𝒜 are sets of bones and angles defined, respectively, by two or three naturally connected joints. To extract kinematic features, we calculated the *L*_2_-norm of the *n*-th derivative of each element in these sets as follows:

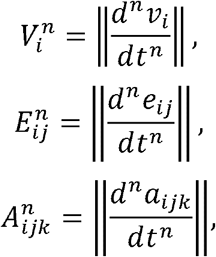

where n = {0,1,2}. Thus, we extract a total of 9 types of kinematic features summarized below:

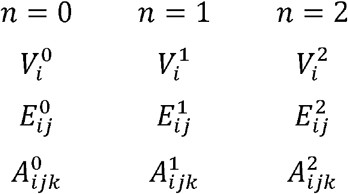

### Action skeleton details

We set our action skeletons to consist of 9 nodes, 8 edges, and 7 angles. To provide an intuitive understanding, we also provide specific nomenclature for defined nodes, edges, and angles, as below:

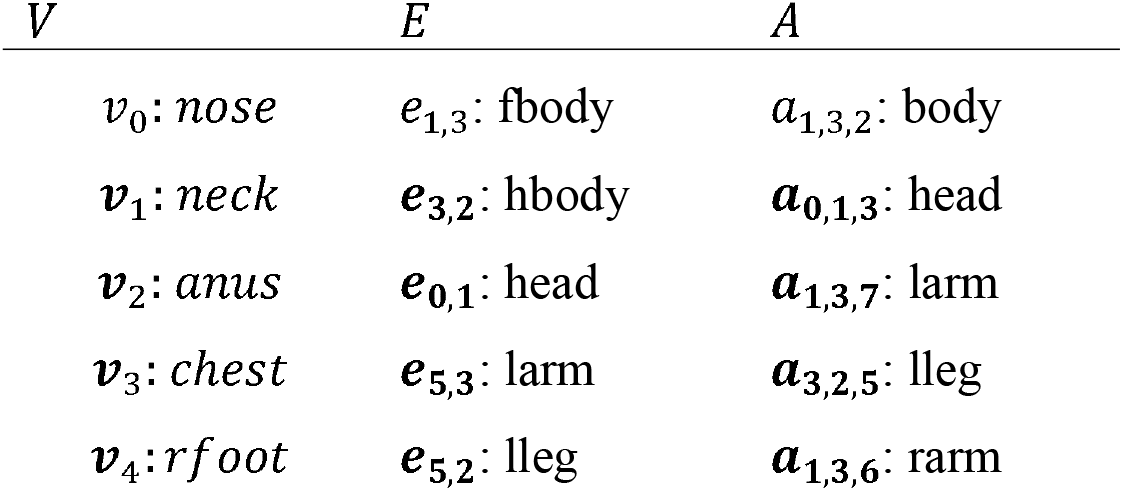
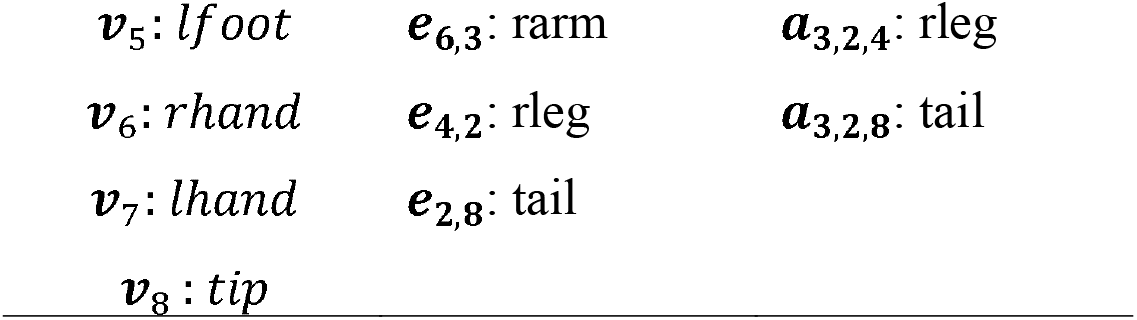

| $V$ | $E$ | $A$ |
| --- | --- | --- |
| $v_0$: *nose* | $e_{1,3}$: fbody | $a_{1,3,2}$: body |
| $v_1$: *neck* | $e_{3,2}$: hbody | $\mathbf{a}_{0,1,3}$: head |
| $v_2$: *anus* | $e_{0,1}$: head | $\mathbf{a}_{1,3,7}$: larm |
| $v_3$: *chest* | $e_{5,3}$: larm | $\mathbf{a}_{3,2,5}$: lleg |
| $v_4$: *rfoot* | $e_{5,2}$: lleg | $\mathbf{a}_{1,3,6}$: rarm |
| $v_5: lfoot$ | $e_{6,3}: rarm$ | $a_{3,2,4}: rleg$ |
| $v_6: rhand$ | $e_{4,2}: rleg$ | $a_{3,2,8}: tail$ |
| $v_7: lhand$ | $e_{2,8}: tail$ | |
| $v_8: tip$ | | |

### Mapping behavioral states

Our method is inspired by a previous spectrogram-based behavior mapping approach (*14, 18*). Here, we describe the details of the method.

### Continuous Wavelet Transformation

First, we extracted *C* channels of spectrogram via wavelet transformation for given *D* euclidean dimensions and *N* nodes; *D* and *N* were 3 and 9, respectively. We used node information only. We employed the Morlet wavelet, which has the following formulation:

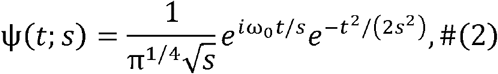

and the continuous wavelet transformation (CWT) of signal is calculated via

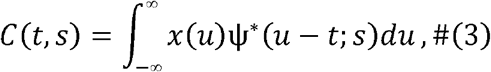

where ω_0_ is the central frequency of the wavelet, *s* is the scale parameter, and *t* is time. ω_0_ determines the dominant frequency content of the Morlet wavelet, while *s* determines the effective time window of the wavelet. The time scale *s* and the fundamental frequency *f* have the following relationship:

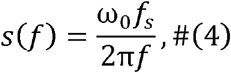

Where *f*_*s*_ is the sampling rate. After the wavelet transformation, the power spectrum *S*(*f*; *t*) (i.e., wavelet spectrogram) is calculated as

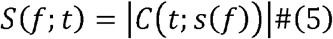

We set ω_0_ = 5 and the frequency range was given as [*f*_min_, *f*_max_], *f*_max_ = *f*_*s*_/2 Hz (Nyquist frequency), and *f*_min_ = *f*_max_ / 10 *HZ*. In our experiment, *f*_s_ was 20 Hz so that *f*_max_ and *f*_min_ were set at 10 Hz and 1 Hz, respectively. The frequency interval between *f*_max_ and *f*_min_ was evenly spaced into 50 segments, i.e., *C* = 50.

Spectrograms were obtained for each node and Euclidean dimension axis. Thus, the preprocessed features were given as multiple spectrograms, and the dimension of the feature at time *t* was *D* × *N* × C. Including the original raw Euclidean coordinates *D* × *N*, the final feature dimension is *D× C×* (*N* + 1) = 3 × 50 × 10 = 1500. The total number of frames per session was 12,000, and the number of sessions was 10. We applied the CWT for each session; after the CWT, we randomly sampled 12,000 data points and applied the principal component analysis (PCA) on this randomly selected dataset due to its computational efficiency. After PCA, we selected the top 100 principal components (PCs) and mapped all preprocessing data (i.e., CWT data) to these 100 PCs. By first reducing the dimensionality of the data using PCA, the computation time for UMAP or t-SNE can be reduced, especially when dealing with high-dimensional datasets.

### Nonlinear embedding methods

Next, we conducted nonlinear dimension reduction into 2-dim from 100 PCs for clustering purposes. This 2-dim embedding space is called *behavior space*. We tested two nonlinear embedding methods t-SNE (*31*) and UMAP (*32*), for creating behavior spaces. *z*_*t*_ denotes a point at time step *t* in the behavior space. The number of neighbors in the UMAP method was set to 50. The perplexity of t-SNE was set at 30.

### Subclustering methods

Applying UMAP or t-SNE to the reduced data can further enhance the clustering results by revealing more structure in the data. With the given set of points in the behavior space or embedding space, we performed Phenograph clustering (*39*) and *k*-Means clustering methods to determine *k* communities or subclusters.

Phenograph clustering was originally designed for high-dimensional single-cell phenotyping. In this method, each node or point in the behavior space is connected to its κ nearest neighbors in the behavior space. Thereby, a κ-nearest neighbor graph is created. After that, Phenograph uses modularity optimization (e.g., Louvain or Leiden) to partition the graph into *k* communities or subclusters of nodes that are more similar to each other than nodes in other subclusters. We utilized the Leiden method, which produces results more quickly than Louvain (*40*). Modularity of the partitioned graph is maximized when nodes are more densely connected within the same subcluster than to other nodes in different subclusters. If transitions between nodes are more likely to happen when the two nodes are closely located in the embedding space than when they are more distantly separated, the trajectory in the behavior space would indicate staying longer in the same community. We call this community or subcluster behavioral state *S*(*t*); the transition matrix **T**(*n*) is defined on this behavioral state, where *n* is the number of steps between two states. We set κ to 30 in the Phenograph algorithm. The Phenograph algorithm requires no predefined number of communities *k* to automatically find the optimal *k* for the modularity of the subdivided graph.

In contrast, the objective of *k*-Means clustering is to divide the entire dataset into *k* clusters such that the sum of squared distances between each point and the cluster center is minimized. In this method, modularity is not taken into account, and *k* must be predetermined.

### Superclustering method

After the subclustering process, we employed the predictive information bottleneck method for obtaining superclusters. Berman et al. (*18*) showed that 1) the transition matrix of fly behavioral states has hierarchical structures and 2) a sequence of behavioral states has a longer time scale than the Markovian assumption, i.e., history of behavioral states affects the next behavior transition. Based on this observation, they proposed dividing behavioral states into *superclusters* that maintain a significant amount of the information regarding future actions (predictive information) that is provided by the current behavioral state.

In a Markovian system, the mutual information between two behavioral states in time *t*_1_ and *t*_2_, *I*(*S*(*t*_1_), *S*(*t*_2_))decays exponentially for large separation |*t*_1_ − *t*_2_| and a characteristic decay time is

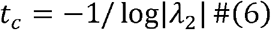

Where λ_2_ is the second largest eigenvalue of T(n = 1). We grouped our behavioral states S(t) into groups Z such that maintains the information about the future state after 2t_c_ transition steps (*18*). In detail, we maximized the following equation:

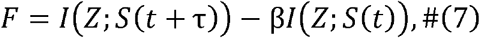

where τ is 2*t*_*c*_, the first term increases the information about the state after τ transitions in the future, while the second term reduces information from the past; β is a parameter that controls the predictive power of the supercluster Z. The deterministic information bottleneck (DIB) algorithm was utilized to optimize this equation, and the optimization procedure followed was the same as in (*18*).

Our model is a sequential combination of CWT, PCA, UMAP (or t-SNE), Phenograph (or *k*-Means), and DIB. For the new dataset, we used a pretrained model for behavioral state mapping. Note that for new embedding 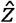,we used the nearest neighbor on the training dataset with previously detected subclusters or communities in the Phenograph stage.

### Temporal connectivity analysis of behavioral state embeddings

To evaluate the temporal connectivity of non-linear state embeddings, we generated discrete behavioral state categories by conducting *k*-Means clustering while increasing *k*. Next, we evaluated the proximal transition index using an inter-cluster transition probability matrix and cluster centroids.

### Comparison of superclusters and human-annotated categories

We used two evaluation metrics: Adjusted Rand Index (ARI) (*41, 42*) and Normalized Mutual Information (NMI) (*43*). The NMI ranges between 0 and 1, and the ARI ranges between - 1 and 1. For all metrics, higher is better, and all of them can accommodate different numbers of classes between the superclusters and the human-annotated categories.

### Manual annotation for behavior categories

Annotation of behavior was performed manually by trained researchers using a custom-built GUI interface. Researchers were instructed to annotate each video frame with one of four predefined categories of behavior: walking, grooming, rearing, and standing. “walking” indicates when the mouse shows movement, specifically with changes in the paw positions. “grooming” indicates when the mouse strokes body parts using their forepaw(s) or licks their body. “rearing” is when the mouse puts weight on its hind legs and lifts its forepaws off the ground. “standing” is when all of the mouse’s paws are on the ground and it is standing still. “na” is the abbreviation for *not annotated*.

## Acknowledgments

The authors would like to thank D. G. Kim and J. J. Park from ACNTNOVA for their helpful discussions and experimental support on the AVATAR system.

## Author contributions

Conceptualization: JK, SPK; Data curation: JK, SPK, JHJ, SHK; Methodology: JK, DKK; Investigation: JK, SPK; Visualization: JK, SPK; Project administration: JK, CJL; Supervision: CJL, MYC; Software: SPK; Writing – original draft: JK, SPK, DKK; Writing – review & editing: CJL, MYC; J.K. and S.P.K. contributed equally to this study.

## Data and materials availability

The data and code used in this study are available at the following GitHub repository: https://github.com/jeakwon/subtle. A demo site showcasing the application of the code is available at https://ibs.re.kr/subtle. Other materials, and codes used in the study are available upon request. All data and code are released under the GNU General Public License v3.0. The authors declare that the data and code support the findings of this study.

## List of Supplementary Materials

Figs. S1 to S7

Table S1

