## Supplementary Figure for "SUBTLE: An unsupervised platform with temporal link embedding that maps animal behavior"

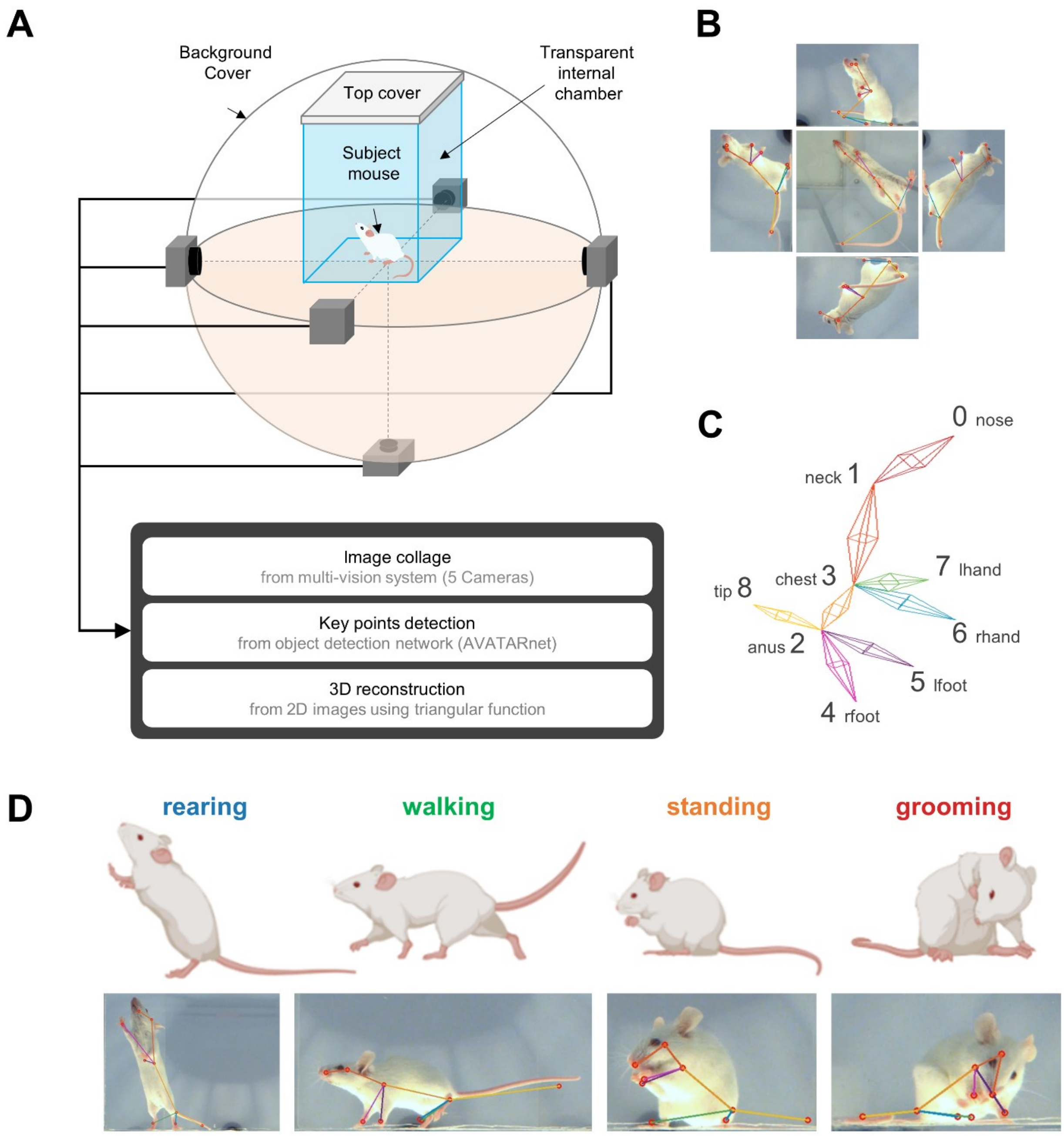


**Fig. S1. 3D action skeleton acquisition system.** (**A**) Action skeleton extraction using 3D pose estimation system. Illustration of recording environment and data extraction steps in the AVATAR system (left) (**B**) Captured snapshots of the subject mouse from 5-way cameras. (**C**) Reconstructed representative action skeletons of various actions with joint numbers and names. (**D**) Example reconstructed 3D skeletons are depicted for 4 behavior categories.

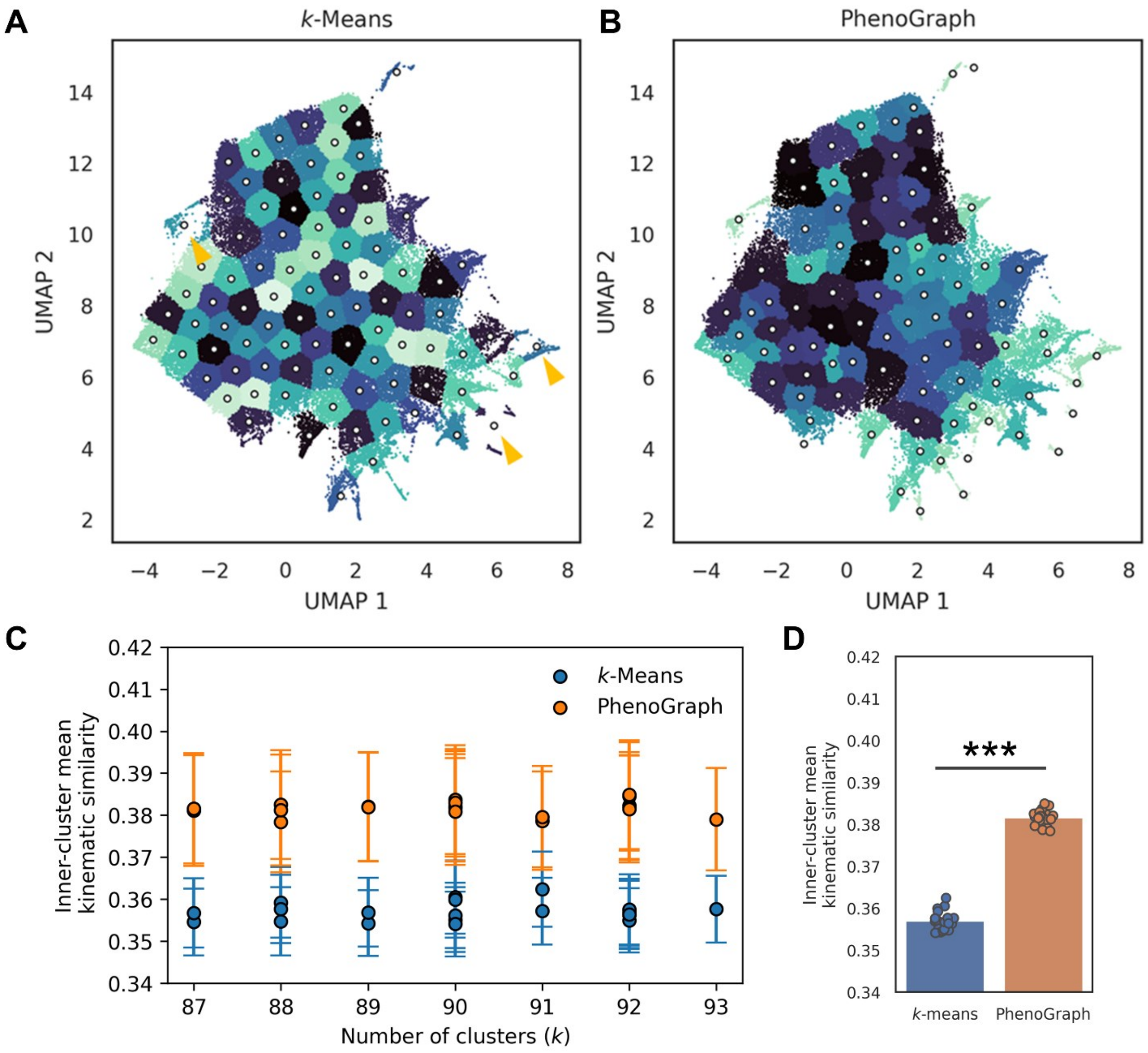


**Fig. S2. Phenograph clustering outperforms *k*-Means clustering for subcluster generation for SUBTLE.** (**A**) Subclusters detected by *k*-Means clustering. Yellow arrows indicate the clusters which contain more than two visually isolated chunks. (**B**) Subclusters detected by Phenograph clustering. (**C**) Comparison of inner-cluster mean kinematic similarity between *k*-Means clustering and Phenograph clustering with 20 trials. (**D**) Summary bargraph of inner-cluster mean kinematic similarity. ($p < 0.001$, Welch's t-test)

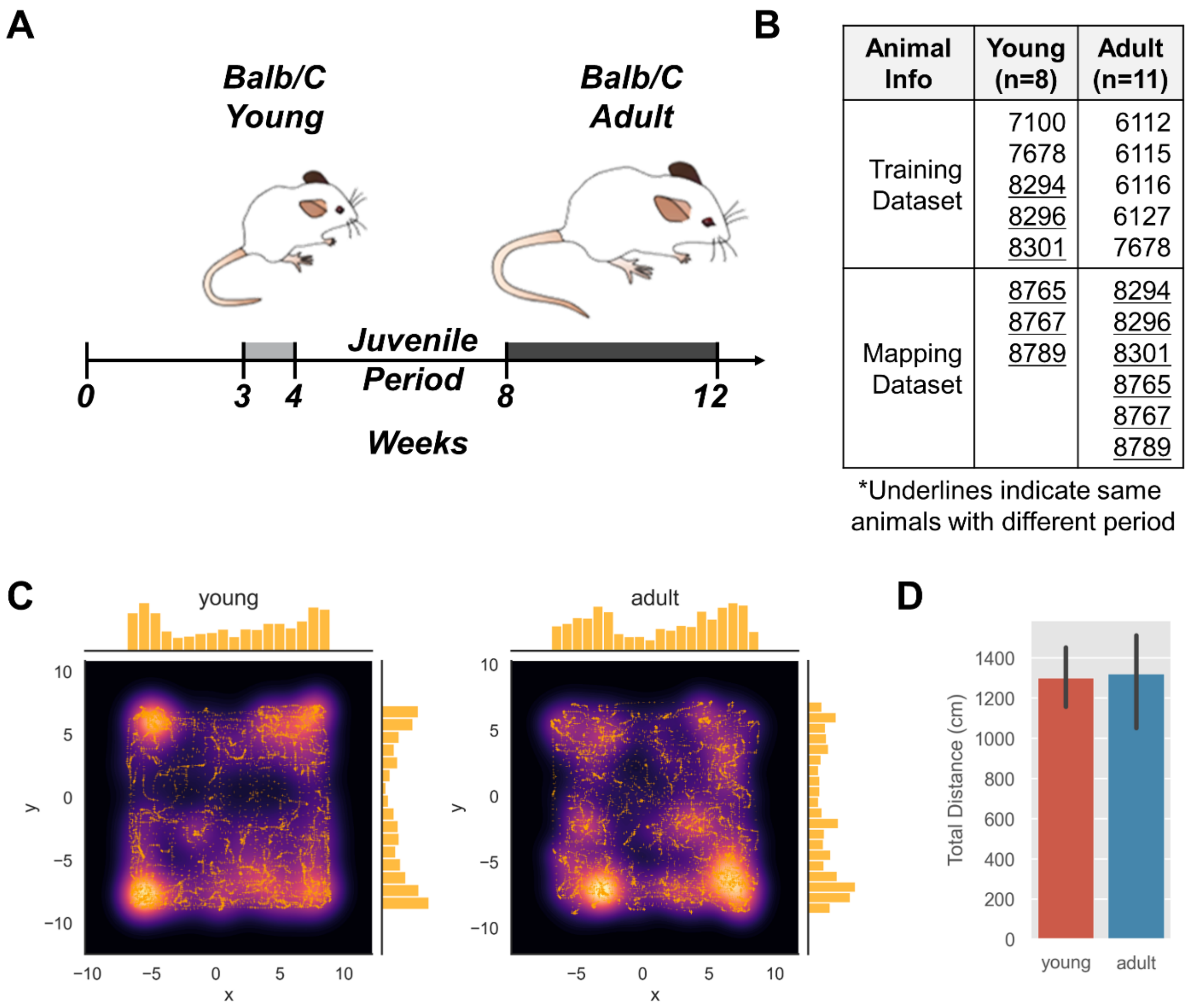


| Animal Info | Young (n=8) | Adult (n=11) |
|---|---|---|
| Training Dataset | 7100<br>7678<br>8294<br>8296<br>8301 | 6112<br>6115<br>6116<br>6127<br>7678 |
| Mapping Dataset | 8765<br>8767<br>8789 | 8294<br>8296<br>8301<br>8765<br>8767<br>8789 |

*Underlines indicate same animals with different period

**Fig. S3. Recorded animal information.** (**A**) Strain and age information of mice. (**B**) Animal ID information for Training and Mapping dataset. (**C**) Example 2D trajectory heatmap of young and adult mice. (**D**) Total distance comparison between young and adult mice (p-value =0.91).

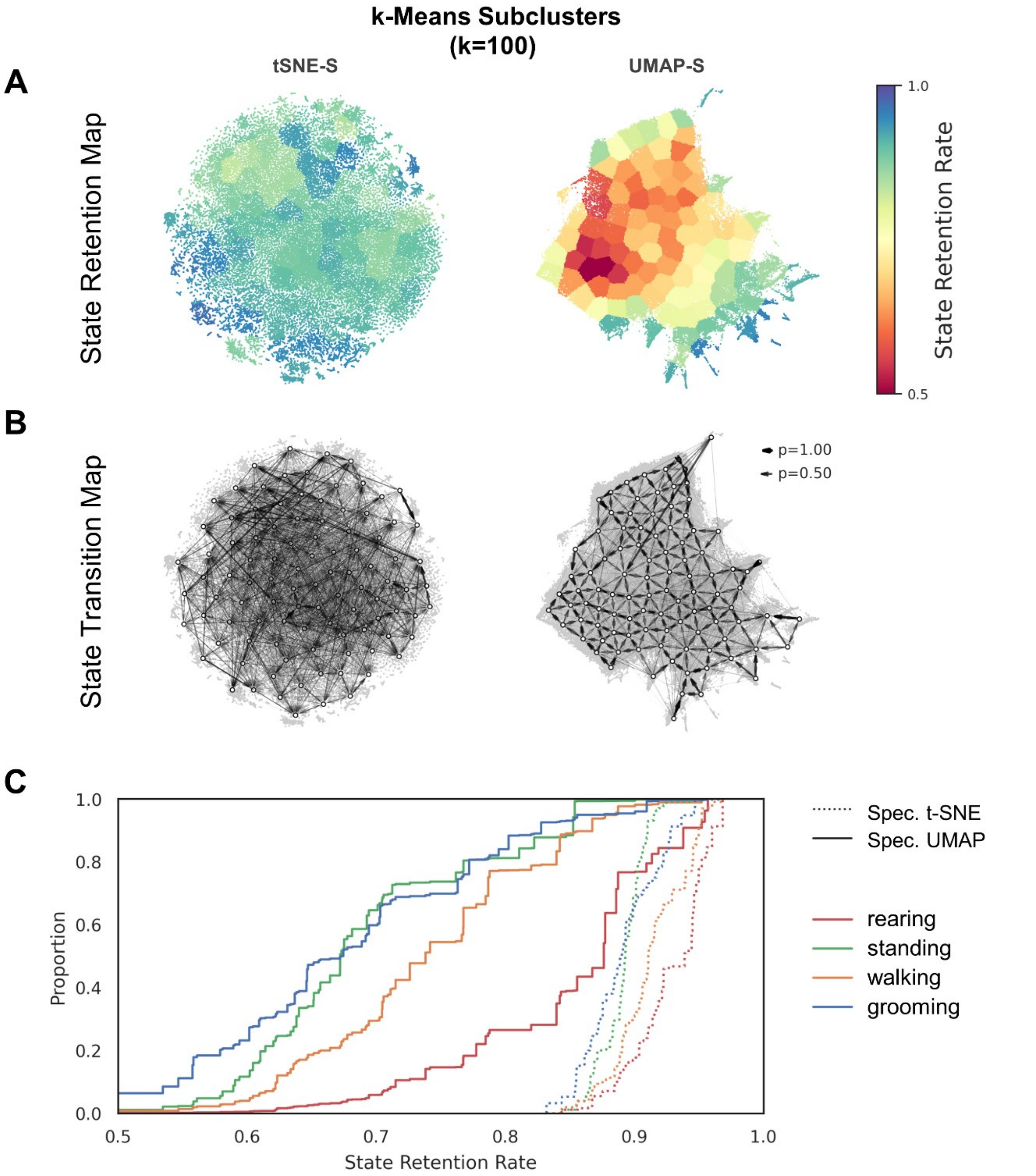


**Fig. S4. State-transition and retention map of subclusters constructed by $k$-Means clustering.** (**A**) State retention map of $k$-Means subclusters ($k = 10$) in embeddings of spectrogram-t-SNE and spectrogram-UMAP (**B**) State transition map of Phenograph subclusters in embeddings of spectrogram-t-SNE and spectrogram-UMAP. (**C**) Empirical cumulative distribution function of human-annotated behavioral states against retention rate.

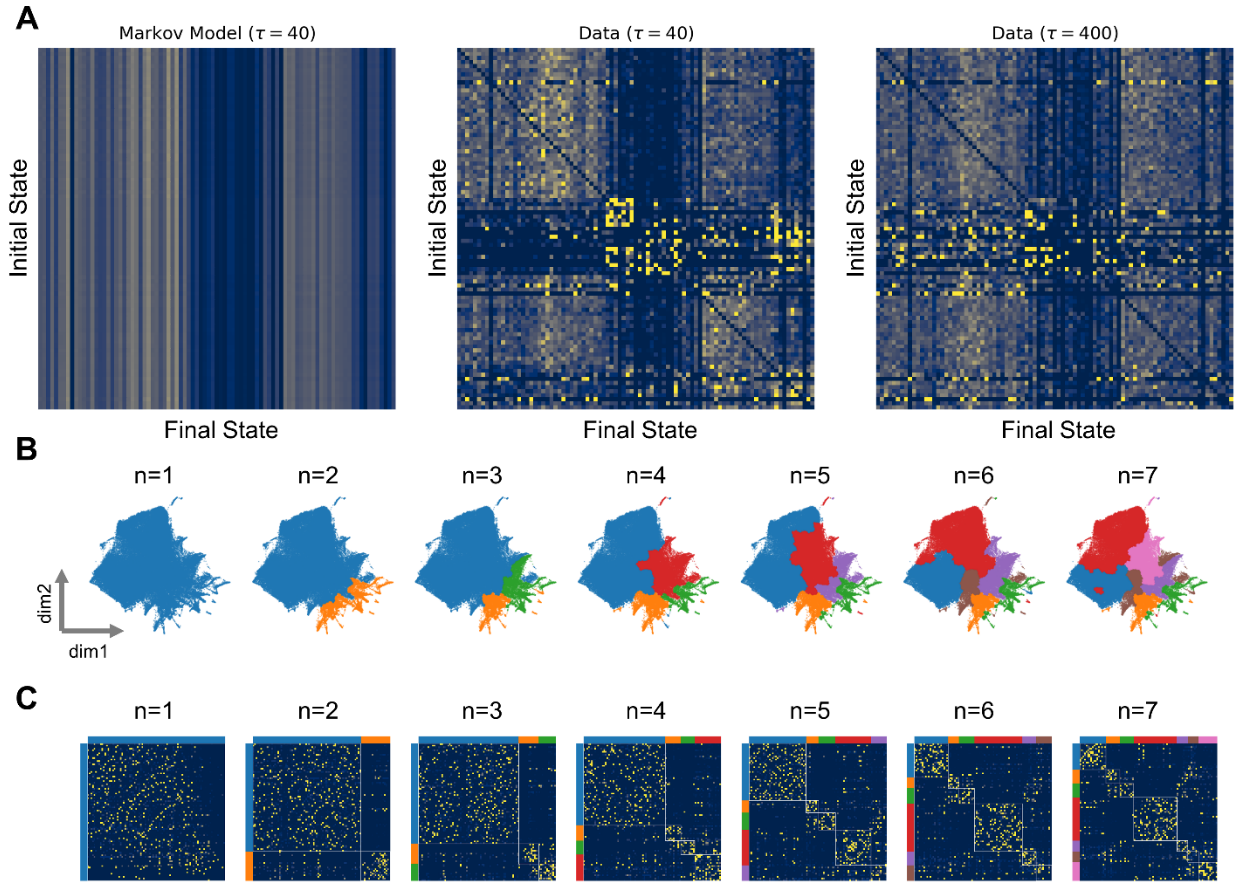


**Fig. S5. Transition probabilities and behavioral modularity for SUBTLE-Pheno-DIB.** (**A**) Markov model transition matrix for $\tau = 40$(left), Transition matrices for $\tau = 40$ (center) and $\tau = 400$ (right). (**B**) Information bottleneck clustering of SUBTLE space for $\tau = 20$ (approximately twice the longest time scale in the Markov model). (**C**) One-step Markov transition probability matrix $\tau = 1$. The 89 subclusters are grouped into $n$ number of superclusters by applying the predictive information bottleneck calculation. White lines denote the cluster boundaries.

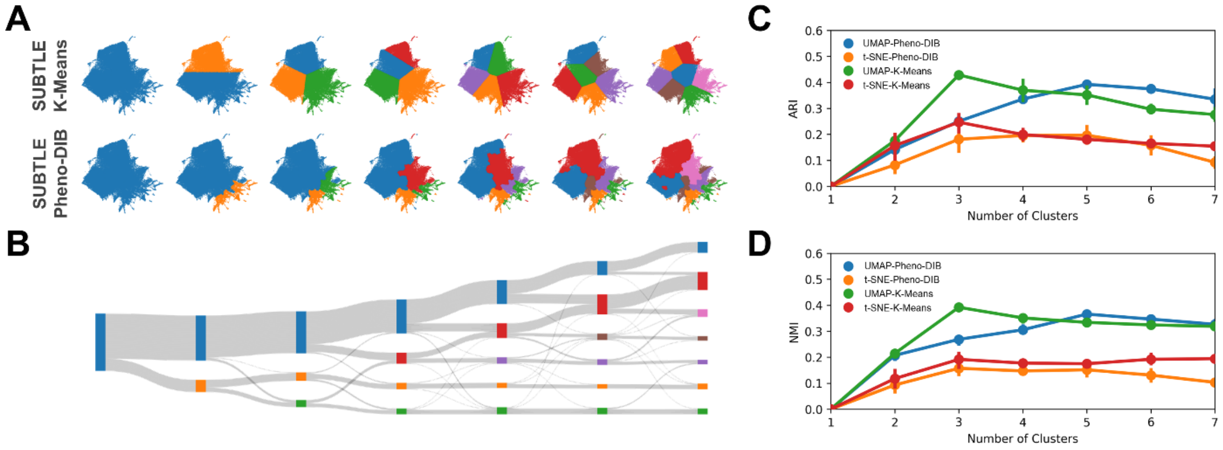


**Fig. S6. Comparison of clustering quality between SUBTLE-Pheno-DIB and SUBTLE-*k*-Means method.** (**A**) Behavior categories depicted on embedding space by SUBTLE-*k*-Means method (top) and SUBTLE-Pheno-DIB (bottom). (**B**) Sankey Diagram of SUBTLE-Pheno-DIB. (**C**) Adjusted rand index scores calculated by comparing with human-annotated categories. (**D**) Normalized mutual information calculated by comparing with human-annotated categories.

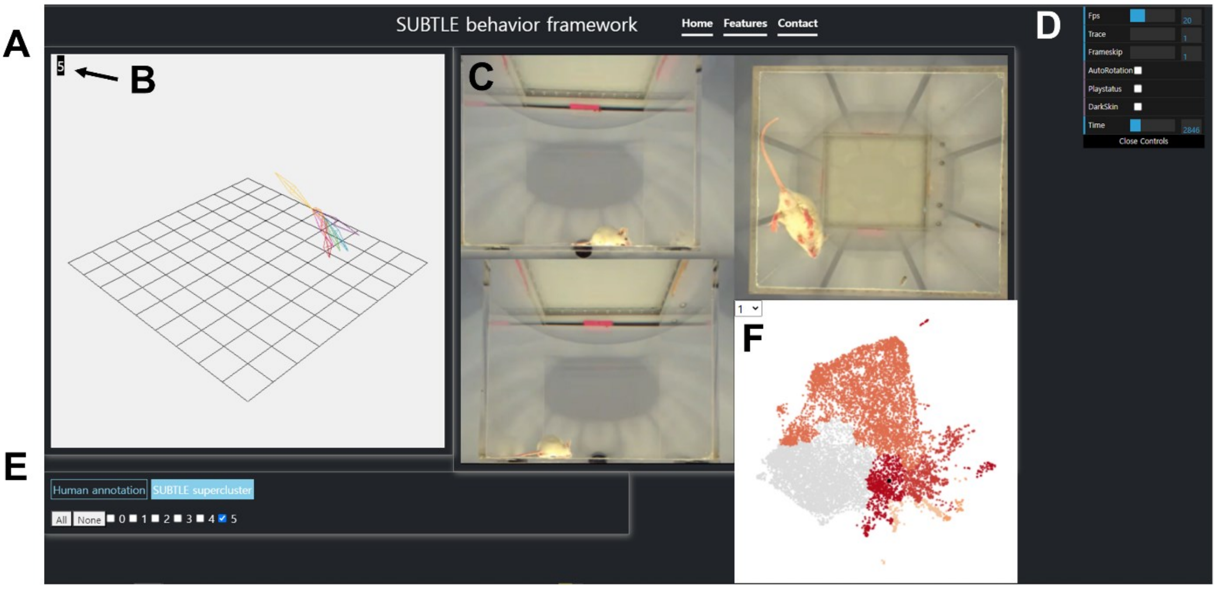


**Fig. S7. SUBTLE visualization framework on the website (https://ibs.re.kr/subtle).** (**A**) 3D action skeleton visualization by Three.js. (**B**) Visualization of current behavior status (human annotation or SUBTLE supercluster-cluster). (**C**) Actual video of mouse behavior taken from AVATAR. (**D**) Controller panel. Fps: Play speed of video; Trace: The number of frames remaining in 3D action skeleton; Frameskip: The number of frames for skipping; AutoRotation: Apply for slow rotation in 3D skeleton; Playstatus: For play or pause the video; Darkskin: Change the color mode (black background) in 3D skeleton; Time: The number of the current frame. (**E**) Annotation filter. (top) “Human annotation” or “SUBTLE supercluster.” (bottom) Checkbox for each annotation label or the number of superclusters. (**F**) The result of behavior mapping after SUBTLE.

| Author (Year)(Ref.) | Method name | Input Features | Nonlinear Reduction | Note |
|---|---|---|---|---|
| Berman et al. (2014)(*14*) | Motion Mapper | C, S | TSNE | Aligned image (Drosophila) |
| Berman et al. (2016)(*18*) | Motion Mapper | C, S | TSNE | Predictive Information Bottleneck Hierarchical clustering |
| Todd et al. (2017)(*35*) | Wavelet-tSNE | S | TSNE | Comparison study PCA vs tSNE |
| Cande et al. (2018)(*19*) | Motion Mapper | C, S | TSNE | Aligned image (Drosophila) |
| Marques et al. (2018)(*44*) | | K | TSNE | Tail, eye tracking (Zebrafish) |
| Pereira et al. (2019)(*24*) | LEAP | S | TSNE | Key-points (Drosophila) |
| Gunel et al. (2019)(*20*) | DeepFly3D | K, S | TSNE | Key-points (Drosophila) |
| DeAngelis et al. (2019)(*45*) | | C | UMAP | |
| Bala et al. (2020)(*21*) | OpenMonkeyStudio | C | UMAP | No detail embedding method |
| Zimmermann et al. (2020)(*23*) | FreiPose (Deeplabcut) | C, S | TSNE | Key-points |
| Jacob et al. (2020)(*46*) | | C | VAE-SNE | |
| Mearns et al. (2020)(*47*) | | D | Isomap | Tail, eye tracking (Zebrafish) |
| Hsu et al. (2021)(*9*) | B-SOiD | K | UMAP | Key-points |
| Huang et al. (2021)(*6*) | Behavior Atlas | K, D | UMAP | Key-points |
| Marshall et al. (2021)(*5*) | CAPTURE | C, K, S | TSNE | Key-points |
| Dunn et al. (2021)(*7*) | DANNCE | C, K, S | TSNE | Key-points |
| Willmore et al. (2022)(*48*) | | C, S | TSNE | k-means clustering 12 behavior features |
| Luxem et al. (2022)(*12*) | VAME | C | Recurrent–VAE- | Deep learning |

**Supplementary Table 1. Summary for various embedding methods used in previous methods** C, raw coordinates (including images); K, kinematic features; S, wavelet spectrograms, D, dynamic time warping.
